# A New Comprehensive Catalog of the Human Virome Reveals Hidden Associations with Chronic Diseases

**DOI:** 10.1101/2020.11.01.363820

**Authors:** Michael J. Tisza, Christopher B. Buck

**Affiliations:** Lab of Cellular Oncology, NCI, NIH, Bethesda, MD 20892-4263

## Abstract

While there have been remarkable strides in microbiome research, the viral component of the microbiome has generally presented a more challenging target than the bacteriome. This is despite the fact that many thousands of shotgun sequencing runs from human metagenomic samples exist in public databases and all of them encompass large amounts of viral sequences. The lack of a comprehensive database for human-associated viruses, along with inadequate methods for high-throughput identification of highly divergent viruses in metagenomic data, has historically stymied efforts to characterize virus sequences in a comprehensive way. In this study, a new high-specificity and high-sensitivity bioinformatic tool, Cenote-Taker 2, was applied to thousands of human metagenome datasets, uncovering over 50,000 unique virus operational taxonomic units. Publicly available case-control studies were re-analyzed, and over 1,700 strong virus-disease associations were found.

## Introduction

The human virome is the sum total of all viruses that are intimately associated with people. This includes viruses that directly infect human cells^1^ but mostly consists of viruses infecting resident bacteria, i.e. phages^2^. While the large majority of microbiome studies have focused on the bacteriome, revealing numerous important functions for bacteria in human physiology^3^, information about the human virome has lagged. However, a number of recent studies have begun making inroads into characterizing the virome^4–12^.

Just as human-tropic viruses can have dramatic effects on people, phages are able to dramatically alter bacterial physiology and regulate host population size. A variety of evolutionary dynamics can be at play in the phage/bacterium arena, including Red Queen^10^, arms-race^13^, and piggyback-the-winner^14^ relationships, to name just a few. In the gut, many phages enter a lysogenic or latent state and are retained as integrated or episomal prophages within the host bacterium^15^. In some instances, the prophage can buttress host fitness (at least temporarily) rather than destroying the host cell. Prophages often contain genes that can dramatically alter the phenotype of the bacteria, such as toxins^16^, virulence factors^17^, antibiotic resistance genes^18^, photosystem components^19^, other auxiliary metabolic genes^20^, and CRISPR-Cas systems^21^, along with countless genes of unknown function. Experimental evidence has shown that bacteria infected with particular phages - i.e. “virocells” - are physiologically distinct from cognate bacteria that lack those particular phages^20^.

There have been a few documented cases where phages have been shown to be involved in increased bacterial virulence^16^ or resistance to antibiotics^22^, demonstrating complex roles for phages in human health. In addition, several studies have conducted massively parallel sequencing on virus-enriched samples of human stool, finding differential abundance of some phages in disease conditions^5,23–25^. A major issue encountered by these studies is that there is not yet a comprehensive database of annotated virus genome sequences, and *de novo* prediction of virus sequences from metagenomic assemblies remains a daunting challenge^2^. In one study, only 31% of the assembled sequence data in virion-enriched virome surveys could be identified as recognizably viral^26^. Impressively, another study of twelve individuals was able to recruit over 80% of reads from virus-enriched samples to putative virus contigs^10^. Still, most of the potential viral contigs were unclassifiable sequences, and a large majority of contigs appeared to represent subgenomic fragments under 10kb.

The current study sought to overcome the traditional challenges of sparse viral databases and poor detection of highly divergent viral sequences by using Cenote-Taker 2, a new virus discovery and annotation tool^27^. The pipeline was applied to sequencing data from nearly 6,000 human metagenome samples. Strict criteria identified over 180,000 viral contigs representing 52,570 specific taxa. In most cases, 75-99% of reads from virus-enriched stool datasets could be back-aligned to the Cenote-Taker 2-compiled Human Virome Database. Furthermore, the curated database allowed read-alignment-based abundance profiling of the virome in human metagenomic datasets, enabling the re-analysis of a panel of existing case-control studies. The re-analysis revealed previously undetected associations between chronic diseases and the abundance of 1,789 specific virus taxa.

## Results

### Characteristics of the Human Virome

Read data was downloaded from NCBI’s Sequence Read Archive (SRA), including data from the Human Microbiome Project^28^ and several other studies (25 Bioprojects in total) involving deep sequencing of human metagenomic samples (Supplemental Table 1)^10,29–42^. These data spanned multiple body sites, including gut (stool), mouth, skin, and vagina. A subset of the projects performed enrichment for viral sequences^43–46^. Almost all of the projects pursued DNA sequencing, but a small number of metatranscriptomic (i.e., ribosome-depleted total RNA) samples were analyzed as well^47^. Read data were binned and assembled by Biosample rather than by individual run to combine read sets from the same individual. A total of 5,996 Biosamples were analyzed.

Cenote-Taker 2^27^ was used to check contigs for two common end-features of complete viral genomes: terminal direct repeats (suggesting a circular or LTR-bounded viral genome) or inverted terminal repeats (ITRs). Sequences with direct terminal repeats were arbitrarily assumed to represent circular DNA genomes. Circularized sequences >1,500 nucleotides (nt) with at least one viral hallmark gene and ITR-containing contigs >4,000 nt with at least one viral hallmark gene were designated as putative viruses. Linear (no discernable end features) contigs >12,000 nt with two or more viral hallmark genes were also kept as putative viruses. Since phages are often integrated into bacterial chromosomes, each linear contig was pruned with the Cenote-Taker 2 prophage pruning module to remove flanking host sequences. The analysis resulted in over 180,000 high-quality putative viral sequences. The sequences were classified into operational taxonomic units (OTUs) by clustering at <0.05 intersequence distance (a proxy for >95% average nucleotide identity) using Mash^48^ (see Methods). A final database of 52,570 sequences representing nonredundant virus OTUs was generated, and this database will herein be referred to as the Cenote Human Virome Database (CHVD, download available at https://zenodo.org/record/4069377).

8,625 virus OTUs consisted of contigs with terminal direct repeats and 113 were ITR-bounded. An additional 2,070 virus OTUs were likely complete proviruses because they were flanked on both sides by host sequences. Since it can be difficult to detect many kinds of viral genome ends using short read assemblies the absence of terminal repeats or flanking chromosomal regions does not necessarily imply that a given contig represents an incomplete viral genome (or genome segment)^49^. It is thus uncertain what proportion of the remaining 41,762 contigs in the CHVD represent complete genomes, as opposed to subgenomic fragments.

Although it is often challenging to obtain very long virus contigs from *de novo* assemblies, 133 virus OTUs over 200 kilobases (kb) were detected in the survey, with the largest being *Siphoviridae* species ctpHQ1, at 501 kb. A total of 38 family- or order-level taxa were observed, and 2,568 sequences could not be classified by Cenote-Taker 2 (Supplemental Fig. 1). It is important to note that virus taxonomy, especially taxonomy of dsDNA phages, is currently in flux ^50–53^, and these taxonomic statistics will likely change as taxonomic groupings are revised. The vast majority of classified sequences represent dsDNA tailed phages in the order *Caudovirales* (Including *Siphoviridae, Podoviridae, Myoviridae, Ackmannviridae, Herelleviridae*, and CrAss-like viruses). Relatively small numbers of known human-tropic viruses were uncovered, including members of families *Adenoviridae, Anelloviridae, Circoviridae, Herpesviridae, Caliciviridae* (Human norovirus), *Papillomaviridae*, and *Polyomaviridae*. Most of the human-tropic viruses mapped to previously reported virus species, but 16 previously undiscovered anelloviruses were detected (download available from: https://zenodo.org/record/4069377). Supplemental Table 2 contains information on each virus, including OTU, hallmark genes, CRISPR hits (see below), and statistical information.

**Figure 1:**
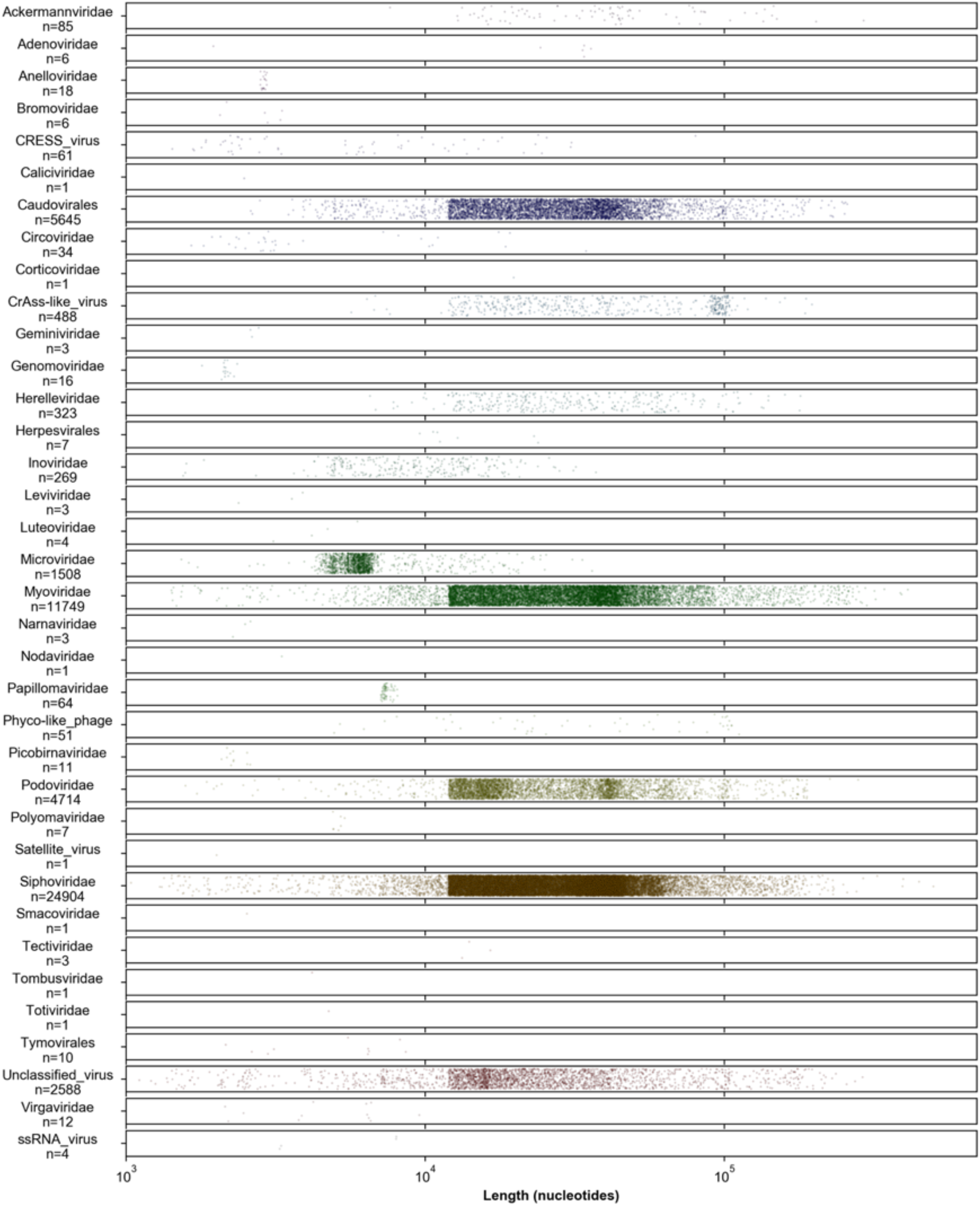
Summary of virus contig taxonomy and length. Each classified contig is represented as dot, with the x-axis position representing contig length. Y-axis values are arbitrary. Contigs smaller than the 12 kb cutoff used for the analysis are either bounded by terminal repeats, or the result of pruning of inferred bacterial chromosomal sequences. Larger (> 3kb) CRESS virus OTUs consist of contigs in a previously reported taxon that combines CRESS-like replication genes with inovirus-like virion genes^54,55^.

Figure 1 presents a graphical summary of observed virus taxa. One taxon, designated “Phyco-like_phage”, is represented by 51 contigs. This is an interesting group of sequences defined by Cenote-Taker 2 as *Phycodnaviridae* due to similarity of the terminase/packaging gene of these viruses to a gene encoded by eukaryotic phycodnaviruses (^~^30% AA similarity). However, most of the inferred virion structural genes that co-occupy these contigs are distantly similar to those of crAss-like phages, not phycodnaviruses, suggesting that they represent phages. This, and the fact that most of the 2,570 “Unclassified” viral sequences have virion hallmark genes corresponding to dsDNA phage models (Supplemental Fig. 1) (Cytoscape network file available from: https://zenodo.org/record/4069377), supports the idea that substantial phage diversity remains unclassified and undescribed.

To evaluate the degree to which the observed CHVD virus OTUs are already represented in public databases, Mash was used to measure roughly intraspecies-level nucleotide sequence similarity to 23,386 genomes from annotated virus species found in GenBank. With a Mash distance threshold of <0.05, 366/52,570 (0.7%) Cenote Human Virome Database viruses had at least one strict cognate sequence in GenBank (Supplemental Table 3). If the Mash distance threshold is relaxed to <0.1, 2,658/52,570 (5.1%) Cenote Human Virome Database viruses have a GenBank cognate (Supplemental Table 3).

Recently, Gregory et al.^12^, published a human Gut Virome Database (GVD) using different virus discovery methods. The Cenote Human Virome Database presented in this manuscript contains sequences from multiple human body sites, so comparisons are not perfect. However, we note that CHVD has 58% more contigs (52,570 vs 33,242) and 181% more sequence information (1.674 gigabases vs 0.596 gigabases) than GVD. The same Mash distance analysis was applied to compare the two datasets. With a strict threshold, 6,832 (13.0%) virus sequences from this study had a cognate in the Gut Virome Database. At the looser threshold, 20,990 (40.0%) sequences had cognates in both databases (Supplemental Table 4).

An important element of the Cenote Taker 2 pipeline is that it provides output files suitable for submission to GenBank. At the time of manuscript submission, the process of depositing all CHVD OTU genome maps into GenBank is ongoing.

**Supplemental Figure 1:**
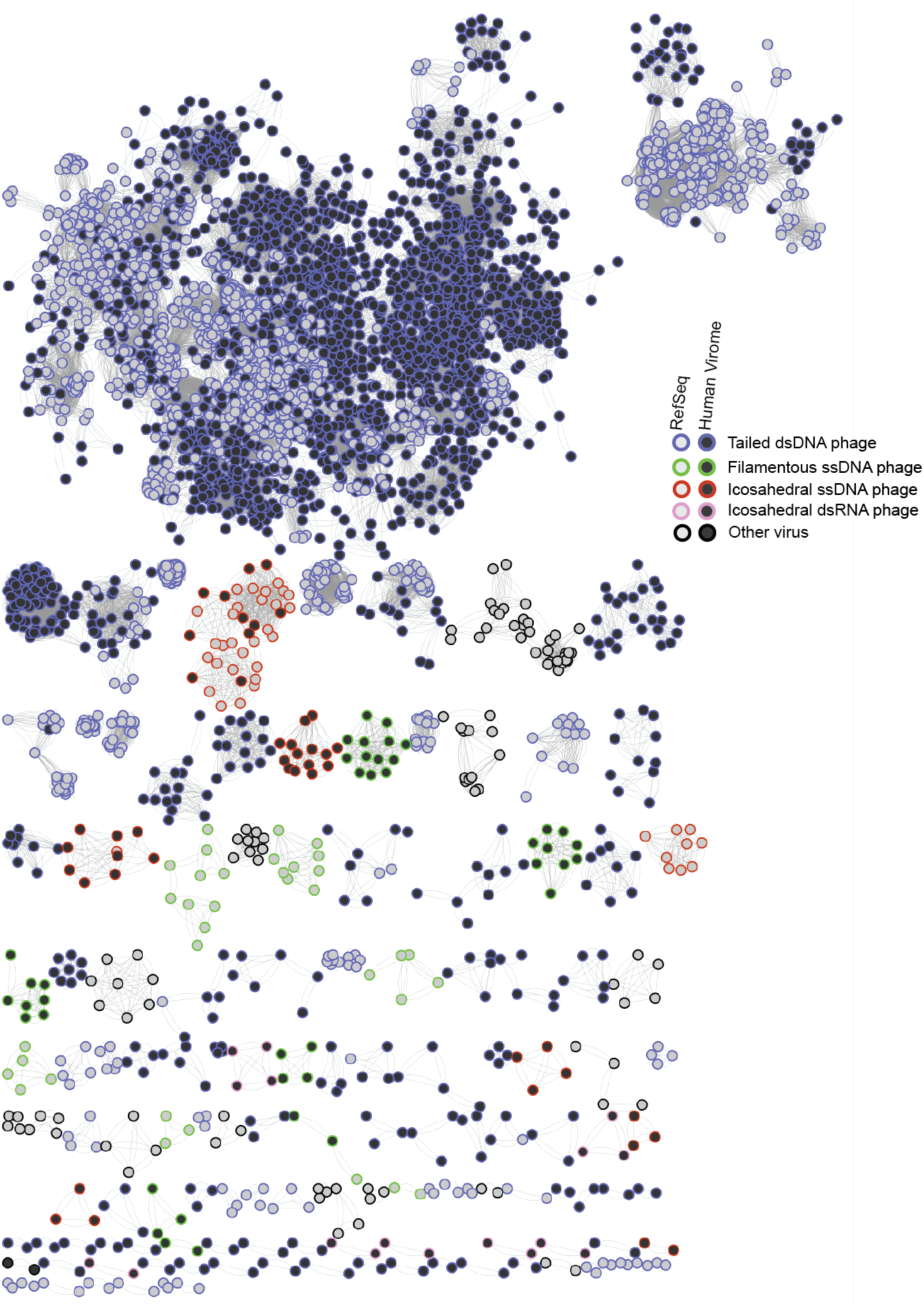
Gene sharing network of RefSeq phages and unclassified OTUs. Vcontact2^52^ was used to make a network of RefSeq bacteriophage genomes and CHVD contigs deemed “unclassified viruses.” Vcontact2 is more sensitive than the taxonomy module of Cenote-Taker 2, as Vcontact2 compares all genes encoded on each contig and Cenote-Taker 2 uses only a single hallmark gene for comparison to its taxonomy database. Edges are drawn between nodes if some proportion of their genes share protein sequence similarity. Only clusters of two or more nodes are displayed. Each sequence cluster was given a feature label, such as “filamentous ssDNA phage,” based on manual inspection of the virion hallmark gene calls made by Cenote-Taker 2 (see Supplemental Table 2) for one or more sequences in the cluster.

### The large majority of reads from well-enriched virome preparations are identifiable

It is unclear how much of the human virome is cataloged by CHVD. One way to address this question is to look at datasets that are physically enriched for viral sequences and determine what fraction of reads in the dataset are identifiable. Reads from 983 human stool samples (representing 11 different studies) that were physically enriched for virions and subjected to nuclease digestion to remove non-encapsidated nucleic acids were mapped against the CHVD (Fig. 2A) (Supplemental Table 5). For most samples with high virus enrichment scores, over 75% of reads were mappable. The result indicates that the database accounts for a large majority of the human virome, at least at a sequence abundance level.

**Figure 2:**
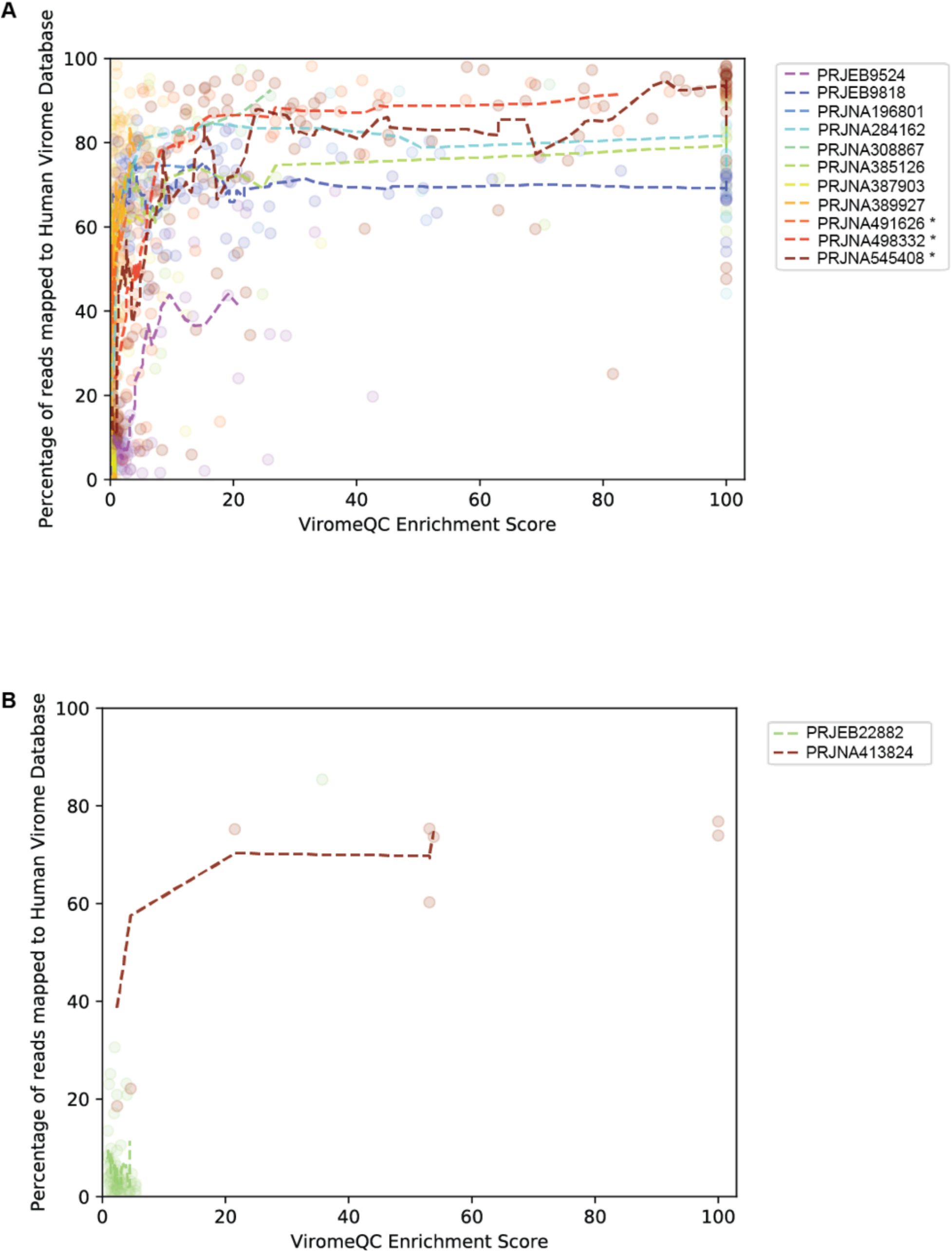
Alignment of reads from “virion-enriched preparations” to the Cenote Human Virome Database. Data from virion-enriched samples are plotted. To measure the degree to which enrichment for viral sequences was achieved, a ViromeQC Enrichment Score^26^ was calculated for each sample (x-axis). The enrichment score is essentially the inverse abundance of known bacterial single-copy marker genes. Dotted lines are moving averages of samples from the same study. Asterisks indicate Bioprojects with data used in the creation of the Cenote Human Virome Database. Panel A shows gut (stool) virome samples, Panel B shows oral (saliva) viromes. A modified database where sequences were clustered at 99% identity instead of 95% identity was used for the index to better capture microdiversity and metaviromic islands^56^ (e.g. intraspecific structural variations consisting of insertions/deletions of gene cassettes, see Methods).

Though well-enriched viromic data were not as available for other body sites, roughly 75% of reads were classifiable in well-enriched oral samples enriched for virus DNA (Fig. 2B) (Supplemental Table 5).

### CRISPR spacer analysis reveals candidate hosts for most phages as well as phage-phage competition networks

Many bacteria encode CRISPR-Cas systems, which contain CRISPR spacer arrays of short (^~^32 nt) sequences copied from and used against invading mobile genetic elements, especially phages^57^. Matching bacterial CRISPR spacers to phage genomes is one way to determine if a bacterial lineage has previously been exposed to a particular phage. Advances in cataloging of CRISPR spacers from bacterial genomes and optimization of phage/host matching pipelines allowed the association of most of the phages discovered in this project to bacterial hosts (http://crispr.genome.ulaval.ca/). Specifically, 36,057 of the 52,570 virus sequences had at least one CRISPR spacer match from a known bacterium or multiple bacteria, with 416,483 total unique spacers matched to Cenote Human Virome Database sequences (Supplemental Table 2). CRISPR spacer density varied dramatically among different bacterial taxa (Fig. 3). For example, members of genus *Bifidobacterium* were confirmed to have relatively large and diverse CRISPR spacer libraries^58^, while *Clostridium, Porphyromonas*, and *Leptotrichia* typically encoded one or only a handful of spacers per phage.

**Figure 3:**
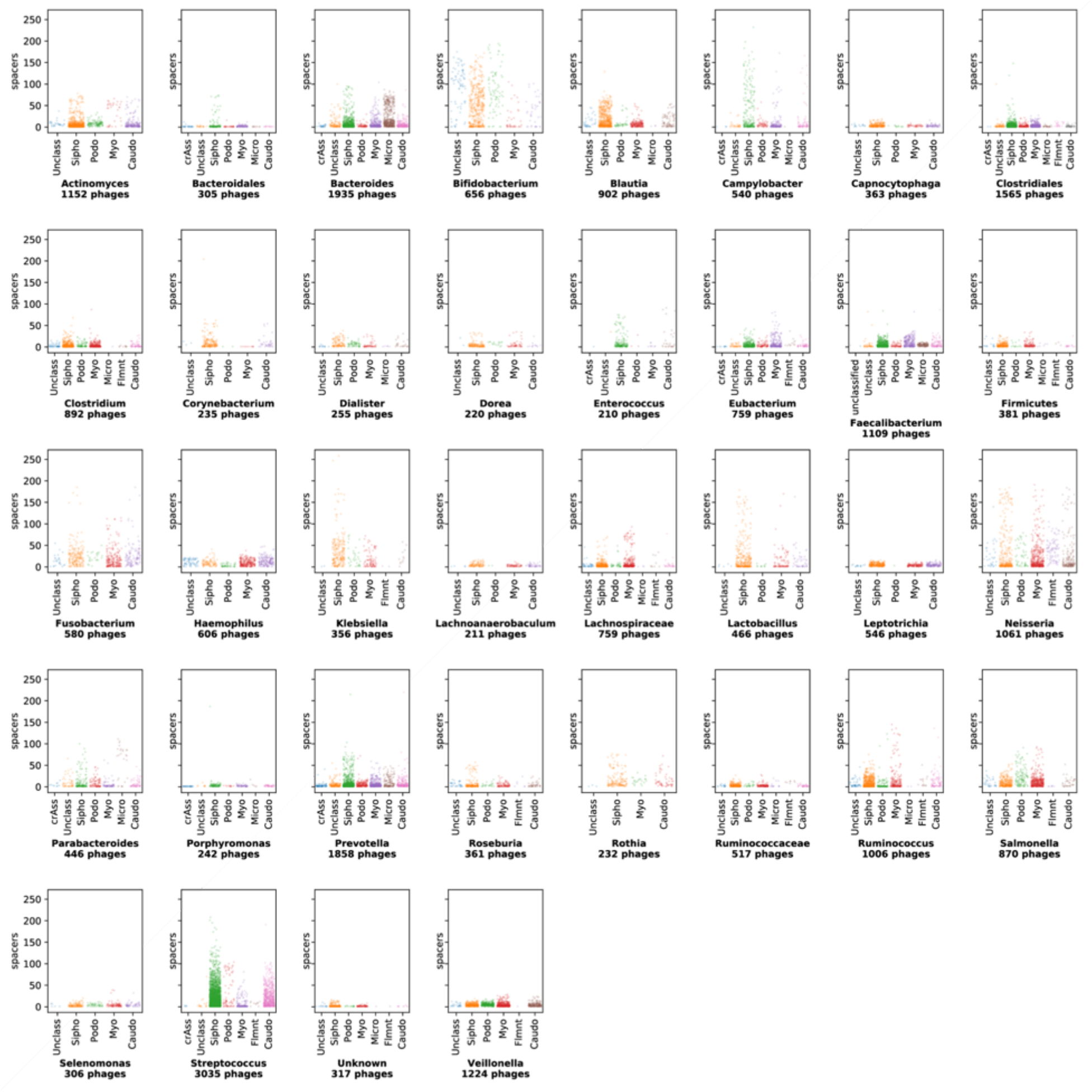
Summary of CRISPR spacer matches to bacterial taxa. Each plot represents data from a different bacterial genus (or higher taxon when genus is not clearly defined, see methods) with CRISPR spacer matches to 200 or more CHVD OTUs. Y-axis values represent the number of unique bacterial CRISPR spacer hits for each virus OTU. Myo = *Myoviridae*, Sipho = *Siphoviridae*, crAss = CrAss-like viruses, Caudo = Other *Caudovirales*, Micro = *Microviridae*, Flmnt = *Inoviridae* and other filamentous phages or CRESS viruses, Unclass = unclassified viruses

CrAss-like viruses seem to be the target of relatively few spacers per virus OTU, despite the fact many representative contigs in CHVD are ^~^100 kb. In contrast, ssDNA phages of family *Microviridae*, despite their small size, seem to be frequently targeted by many unique spacers in *Bacteroides* and *Parabacteroides* CRISPR systems but not CRISPR systems in other bacterial taxa.

Phages themselves can encode CRISPR arrays and some phages have intact and functional CRISPR-Cas systems^21,59^. These CRISPR components can target host defenses as well as other phages competing for the same host^60^. Among phage sequences in the Cenote Human Virome Database, 2,091 CRISPR spacers were detected in arrays from the genomes of 223 phages. Of these, 875 spacers targeted a total of 2,491 other phages, suggesting complex phage-phage competition networks in human metagenomes (Fig. 4) (download cytoscape file from: https://zenodo.org/record/4069377).

**Figure 4:**
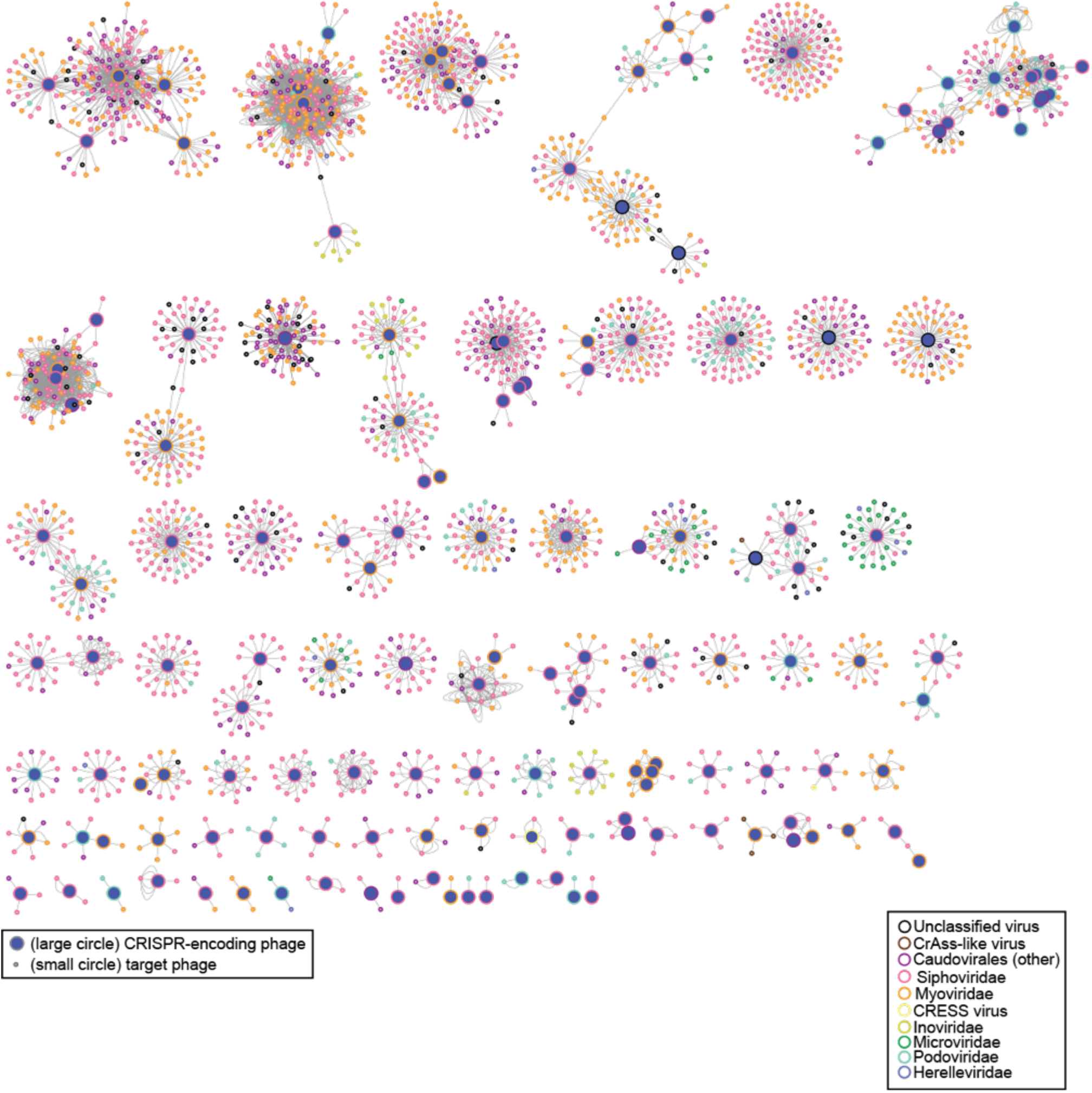
Phage-encoded CRISPR spacers reveal phage interaction networks in the human virome. Network diagram of phage-phage interaction landscape based on CRISPR spacer matches. Each line represents a match of a particular spacer sequence to its target phage.

### The most commonly abundant viruses on several body sites

With this library of viruses and the large sampling effort from the Human Microbiome Project^28,61^, the question of “which viruses are the most commonly abundant” for a given body site can be answered more confidently than was previously possible. It should be noted that the Human Microbiome Project data was collected from healthy Americans between 18 and 40 years of age and the conclusions here may not be generalizable to other populations.

It is often challenging to precisely determine the border between an integrated prophage and host chromosomal sequences without experimental validation. Our preliminary analyses revealed that inclusion of even a few hundred nucleotides of flanking host sequence in a viral OTU can greatly distort abundance measurements because of inadvertent measurement of uninfected host bacterial sequences. We therefore used Cenote-Taker 2 to perform a more stringent analysis in which contigs were trimmed from the first recognized virus hallmark gene through the last virus hallmark gene. We refer to these more stringent units as “virus cores” (download from: https://zenodo.org/record/4069377)

Data were downloaded from SRA and analyzed for hundreds of patients at six body sites (anterior nares, buccal mucosa, posterior fornix, tongue dorsum, supragingival plaque, and gut (stool)). Reads were then aligned to the more stringent virus cores database. As a proxy for the relative abundance of a given virus OTU, the average number of reads per kilobase of virus genome per million reads in the parent dataset (RPKM) was calculated for each sequence, (Fig 5, Supplemental Figs 2–4). Virus prevalence was determined as proportion of samples with >0.1 RPKM. The most commonly abundant virus OTUs were calculated as (mean RPKM ⃗ prevalence). The right panel of Fig 5 shows the inferred host for each of the top 50 most commonly abundant virus OTUs based on CRISPR spacer target information. A majority of the most of the commonly abundant viruses appear to infect members of the common bacterial phylum *Bacteroidetes*, which is generally abundant in the human gut.

**Figure 5:**
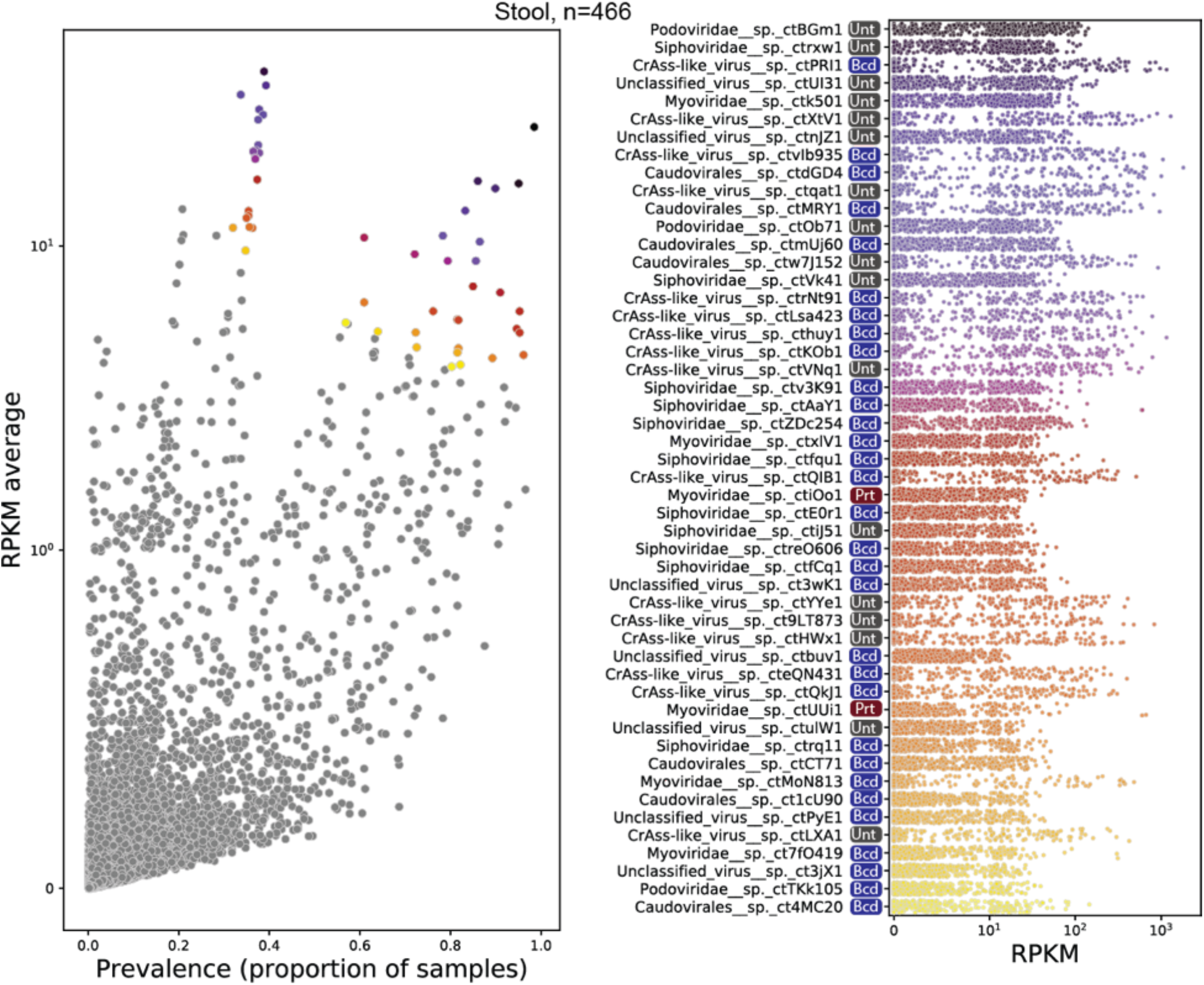
Most common viruses, Stool (Gut) The left panel shows a scatter plot of RPKM (a measure of relative read abundance for a given virus OTU, y-axis) versus prevalence (proportion of samples with >0.1 RPKM, x-axis). For display purposes, the y-axis is a linear scale from 0-1 (10^0^) and log_10_ above 1. The top fifty most commonly abundant virus OTUs (based on the product of coordinates) are colored. Right panel: RPKM values across all samples for the most commonly abundant virus OTUs. Colors of dots in the left panel correspond with the dot strip colors in the right panel. The x-axis is linear from 0-10 (10^1^) and log_10_ above 10. Colored CRISPR target logos: Bcd = *Bacteroidetes*, Prt = *Proteobacteria*, Unt = no CRISPR spacers detected (untargeted).

The data suggest an interesting bifurcation in prevalence of virus OTUs with high abundance (RPKM). Although some virus OTUs, such as *Podoviridae* sp. ctBGm1 and *Myoviridae* sp. ctiOo1, are present in nearly all samples and have an average abundance of >10 RPKM, perhaps representing prophage of ubiquitous bacterial lineages. Others, including all displayed CrAss-like viruses and *Caudovirales* sp. ctdGD4, are absent or low-abundance in most samples but highly abundant in a minority of samples. The latter group could either represent viruses that periodically undergo large replicative bursts, or viruses that constitutively dominate the virome in certain individuals but not others.

**Supplemental Figure 2:**
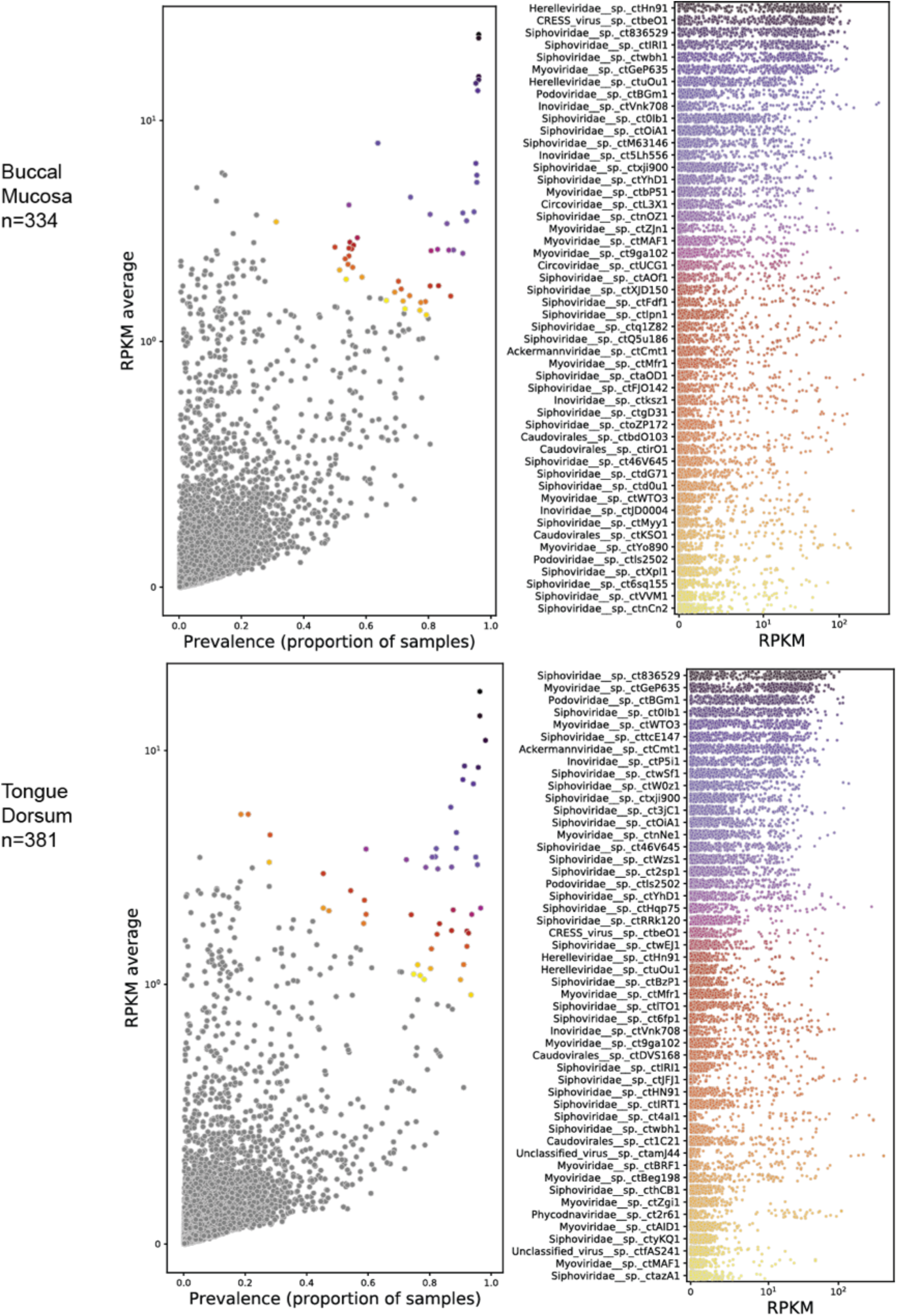
Most common viruses, Buccal Mucosa and Tongue Dorsum. Scatter plots of virus quantification data displayed as described in the legend of Figure 5

**Supplemental Figure 3:**
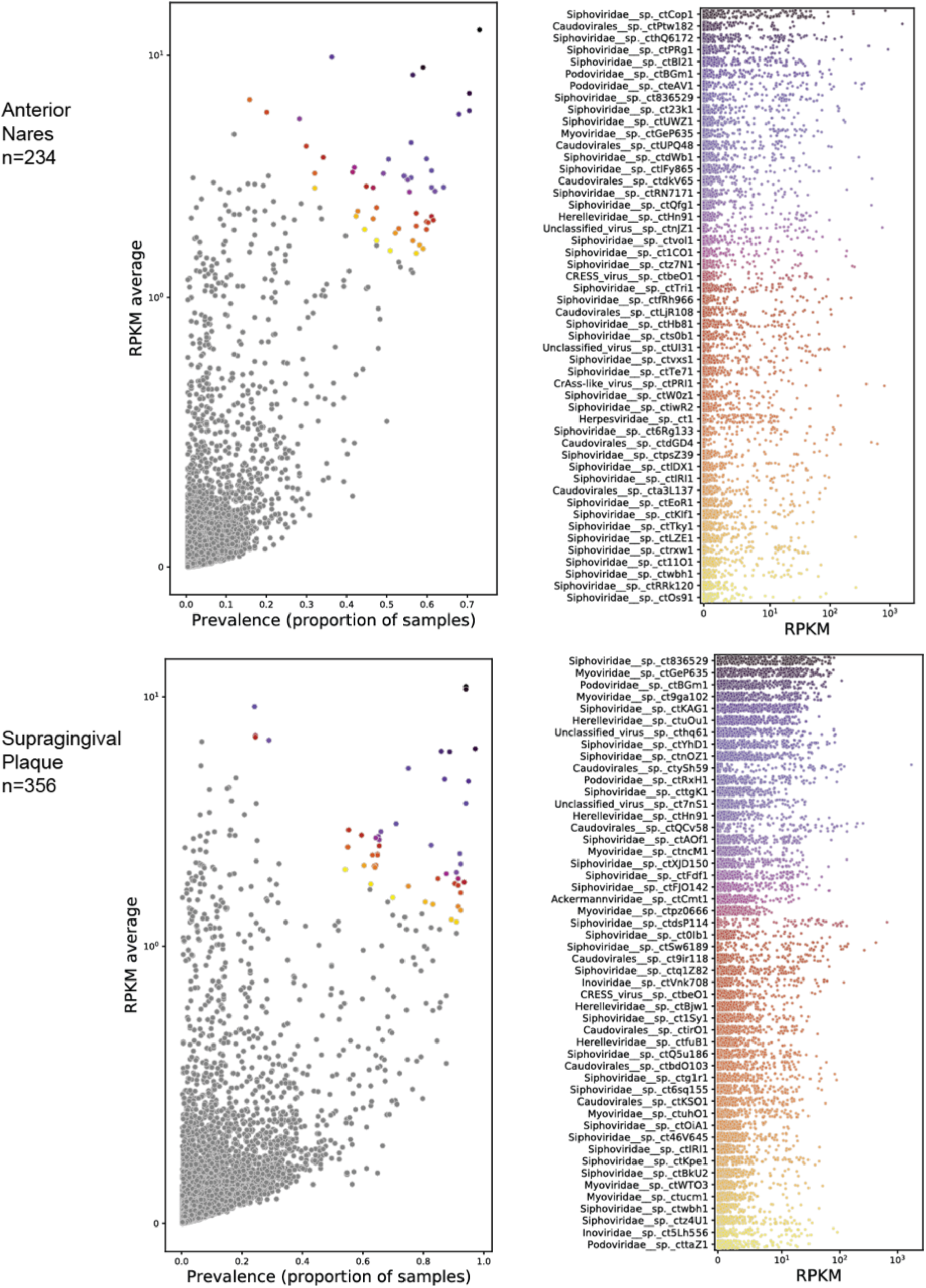
Most common viruses, Anterior Nares and Supragingival Plaque. Scatter plots of virus quantification data displayed as described in the legend of Figure 5

**Supplemental Figure 4:**
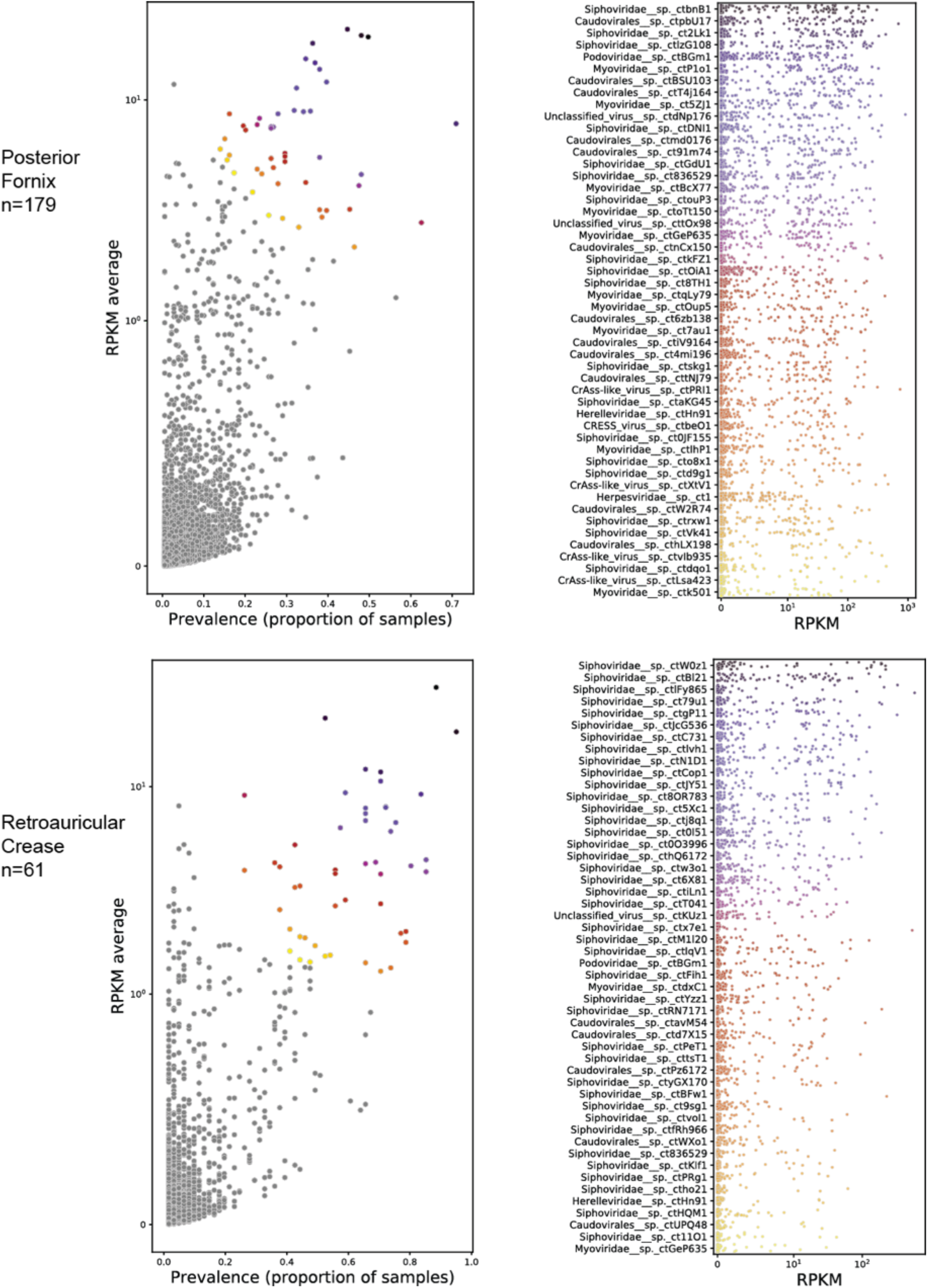
Most common viruses, Posterior Fornix and Retroauricular Crease. Scatter plots of virus quantification data displayed as described in the legend of to Figure 5

### Specific virus OTUs are associated with human disease

A number of prior studies have looked for associations between the virome and human diseases^23–25,46,62–65^. However, these studies were limited by the lack of a comprehensive virus reference database, and nearly all studies examined samples selectively enriched for viral sequences^65^. Virus enrichment methods can be highly variable and can inadvertently remove some viral taxa while failing to significantly select against host sequences^9 26^. Indeed, Gregory et al.^12^ report that studies employing different virus-enrichment protocols to investigate the same disease state (e.g. inflammatory bowel disease) rarely contain the same virus populations in their data. Instead, studies using similar enrichment protocols (regardless of disease state of patients) shared more virus populations. Furthermore, sequences encapsidated within virions may not be the best reflection of the total viral population, especially in human digestive tracts, where many phages are believed to exist primarily in lysogenic (non-lytic) states^66^, and some have been ‘grounded’, losing their ability to independently excise from the host genome^67^. It is possible that the most important phages for human physiology are those that express accessory genes from an integrated provirus state, as opposed to phages that are producing virions. It is thus ideal to examine total DNA (also known as whole genome shotgun, WGS) sequencing, which can detect all DNA virus genomes.

Our study reanalyzed publicly available WGS data from twelve large case-control studies analyzing stool and/or saliva^29,35,68–74^. These studies examined Parkinson’s disease, obesity, colon carcinoma, colon adenoma, liver cirrhosis, type 1 diabetes, ankylosing spondylitis, atherosclerosis, type 2 diabetes, hypertension, and non-alcoholic fatty liver disease. The virus cores database was used to compare the abundance of each virus OTU between case and control cohorts. Figure 6 shows an analysis of case-control comparisons of Parkinson’s disease (Fig. 6 A-C) and obesity (Fig. 6 D-F). RPKM was used to measure virus OTU abundance in each sample, and Wilcoxon rank-sum tests with 100 bootstraps were conducted for each comparison to estimate the p value (Fig. 6A,D “Virome”, Supplemental Figs. 5,6). Significant virus OTUs were determined by a false discovery rate < 1% (see Methods). All analyses compared associations between the virome and the “bacteriome,” measuring the bacteriome in terms of bacterial OTUs (i.e. species-level single-copy bacterial marker gene abundance) using IGGsearch^75^ (Fig 6A,D “Bacteriome”, Supplemental Figs. 5,6). A higher number of statistically significant taxa were found for the virome than the bacteriome in seven studies. The five other studies analyzed yielded no significant OTUs for either the virome or the bacteriome (Fig. 6, Supplemental Figs. 5,6). P values for each virus OTU detected in each study are documented in Supplemental Table 2. Furthermore, random forest classifiers, trained on either all virus OTUs or all bacterial OTUs, were more successful or equally successful, on average, in discriminating healthy and diseased patients using the virome data rather than the bacteriome data in 7/12 case-control populations (Fig 6 B,E, Fig S7,8).

**Figure 6:**
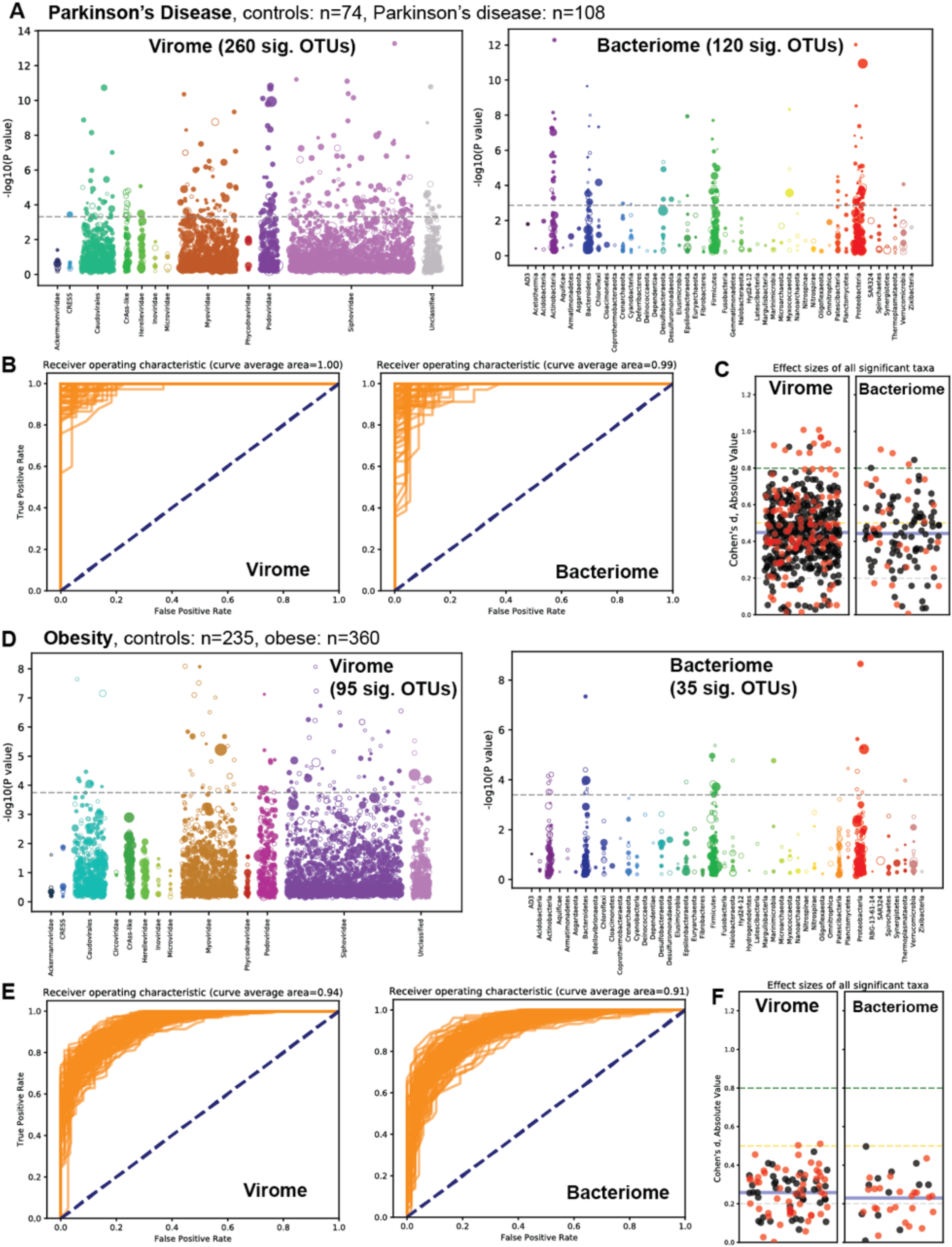
Association of the virome and bacteriome with chronic diseases. (A-C) Analysis of read data from PRJEB17784, a case-control study of stool samples from patients with or without Parkinson’s disease. (A) Virome-wide and bacteriome-wide associations in stool samples from Parkinson’s disease patients (n=74) and healthy controls (n=108) represented as Manhattan plots. Each OTU is represented as a dot along the X-axis, with its y-axis value being the inverse log_10_ p value. The size of each dot corresponds to the median relative abundance of the taxon in the disease cohort. Filled dots represent OTUs found at higher abundance in the diseased state while hollow dots represent decreased abundance in the diseased sate. The dashed grey line represents the false discovery rate < 1% threshold. (B) Receiver operating characteristic plots from 100 differently-seeded random forest classifiers trained on the virome (left) or bacteriome (right). (C) Swarm plots of Cohen’s d effect sizes (absolute value) of OTUs achieving significant p values. Black dots are positive effect size and red dots are negative effect size. The mean of all plotted effect sizes is drawn as a blue line. Small effect size = 0.2 - 0.5; Medium effect size = 0.5 - 0.8; Large effect size = > 0.8^77^. (D-F) Similar analyses of read data from PRJEB4336, a WGS survey of stool samples from obese and non-obese individuals. Plots D, E, and F are laid out in the same manner as plots A, B, and C respectively.

The importance of considering effect size when reporting microbiome associations has become apparent in recent years^76,77^. Therefore, for all virus and bacterial OTUs with significant differences between cases and controls, Cohen’s d effect size is reported for each disease state (Fig. 6C,F, Fig. S7,8).

**Supplemental Figure 5:**
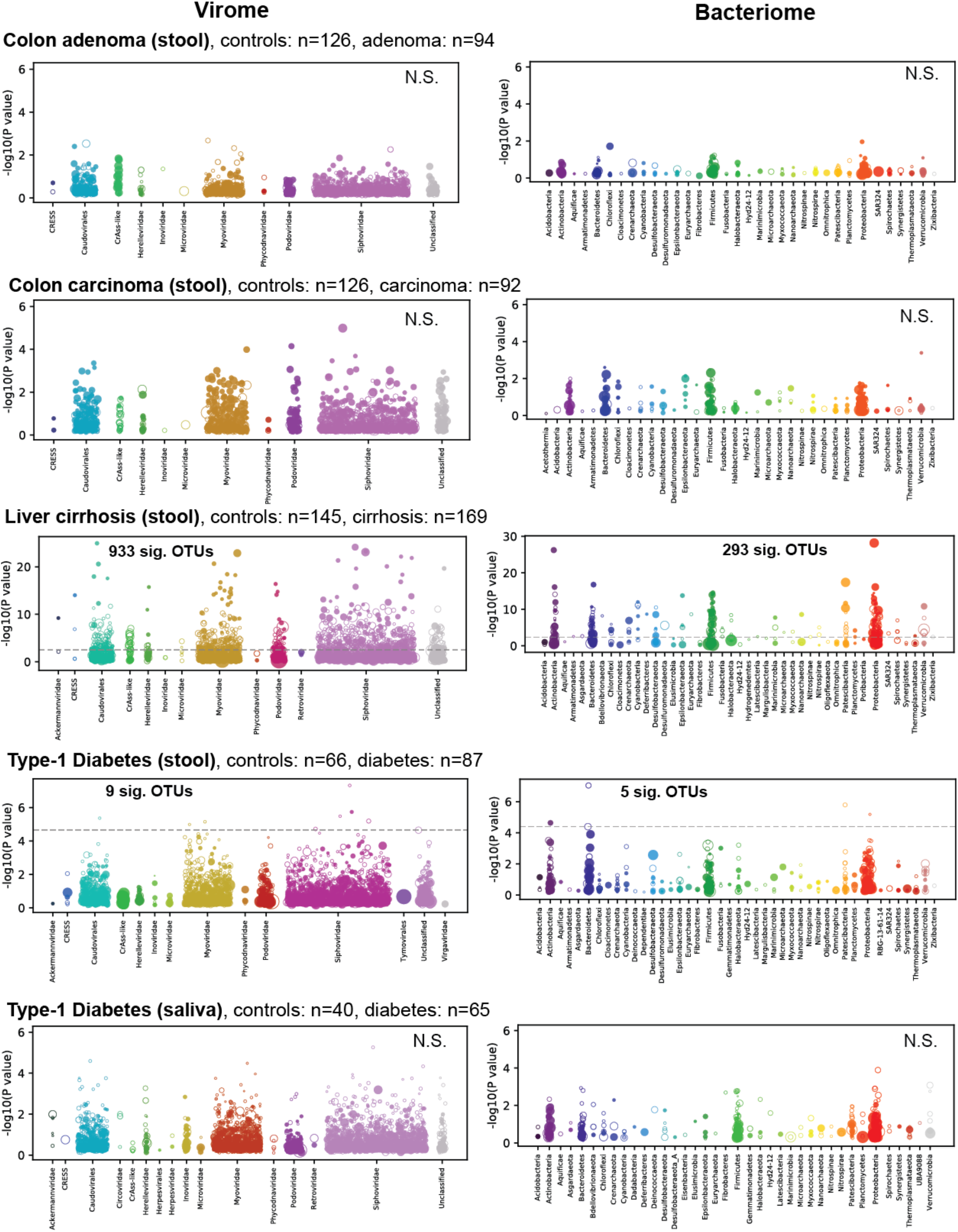
Virome- and Bacteriome-wide associations with additional chronic diseases 1. Manhattan plots for viromes (left) and bacteriomes (right) are shown in the same manner as Figure 6. Accessions: Colon carcinoma and adenoma (PRJEB7774), Liver cirrhosis (PRJEB6337), Type 1 diabetes (PRJNA289586).

**Supplemental Figure 6:**
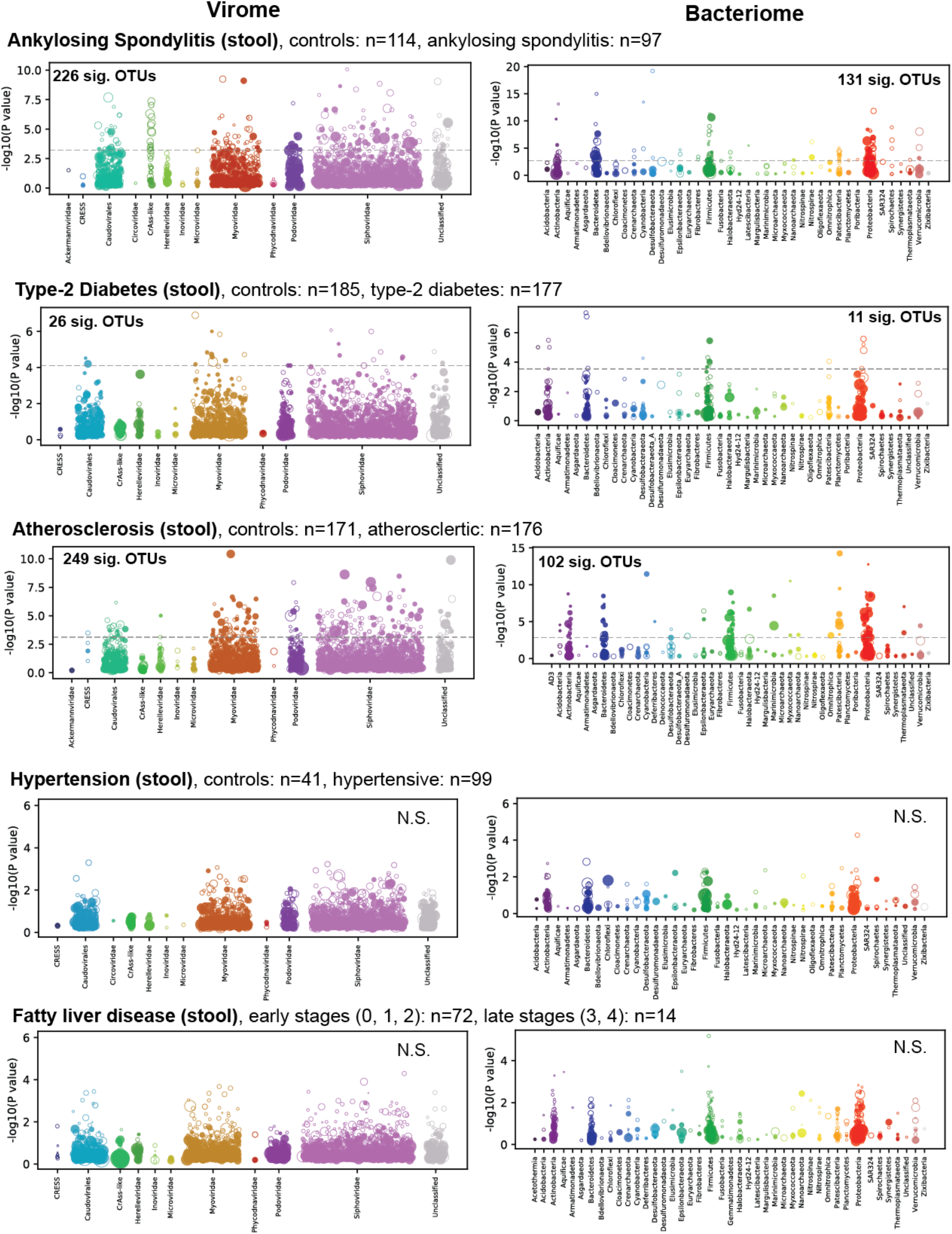
Virome- and Bacteriome-wide associations with additional chronic diseases 2. Manhattan plots for stool viromes (left) and bacteriomes (right) are shown in the same manner as Figure 6. Accessions: Ankylosing spondylitis (PRJNA375935), Type 2 diabetes (PRJNA422434), Atherosclerosis (PRJEB21528), Hypertension (PRJEB13870), Fatty liver disease (PRJNA373901).

**Supplemental Figure 7:**
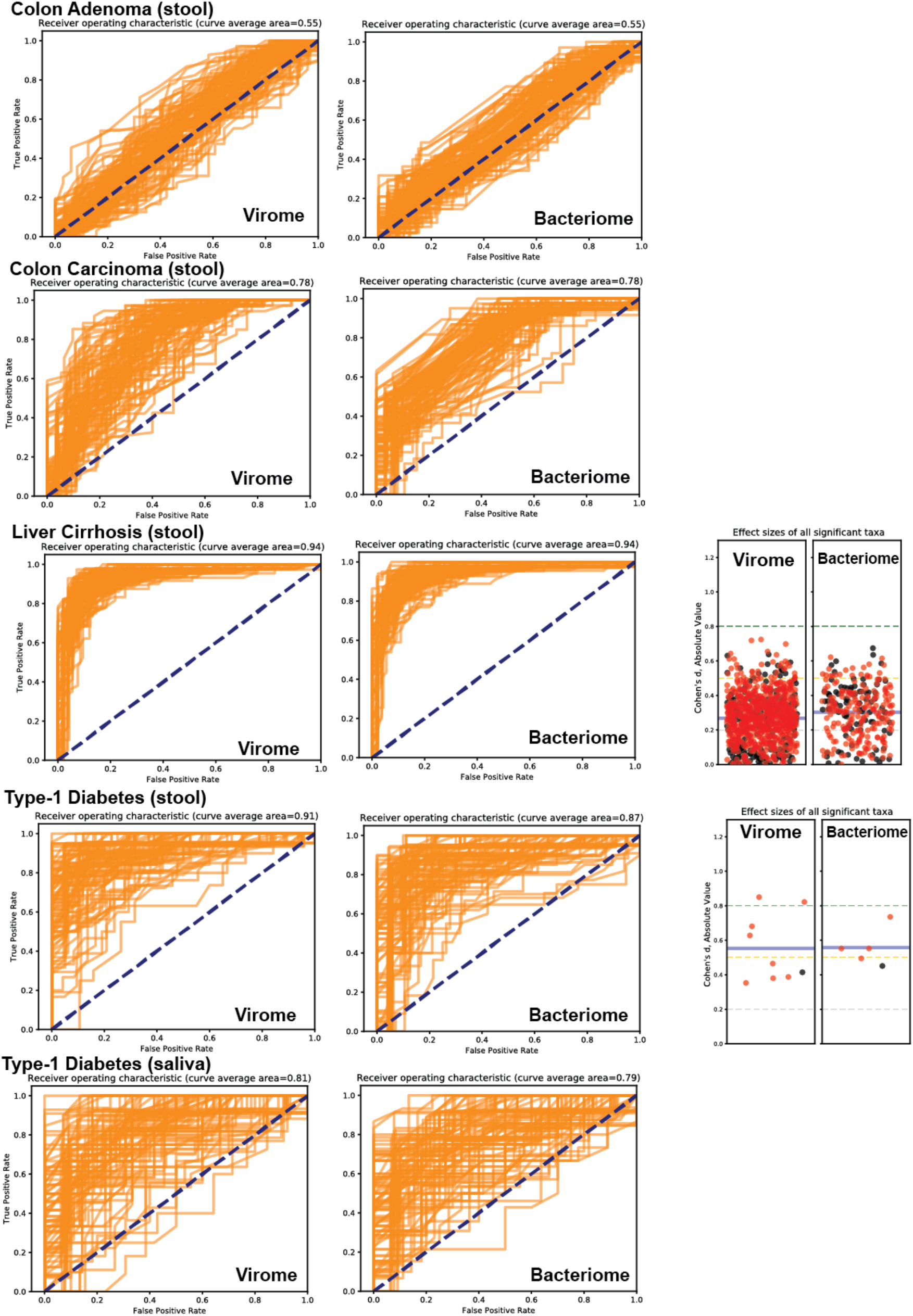
Discriminatory power of viromes and bacteriomes in additional chronic diseases 1. Receiver operating characteristic plots for virome (left panels) and bacteriome (middle panels) data, as described in Figure 6. Summary of effect size data for significant OTUs (right panel), as described in Figure 6.

**Supplemental Figure 8:**
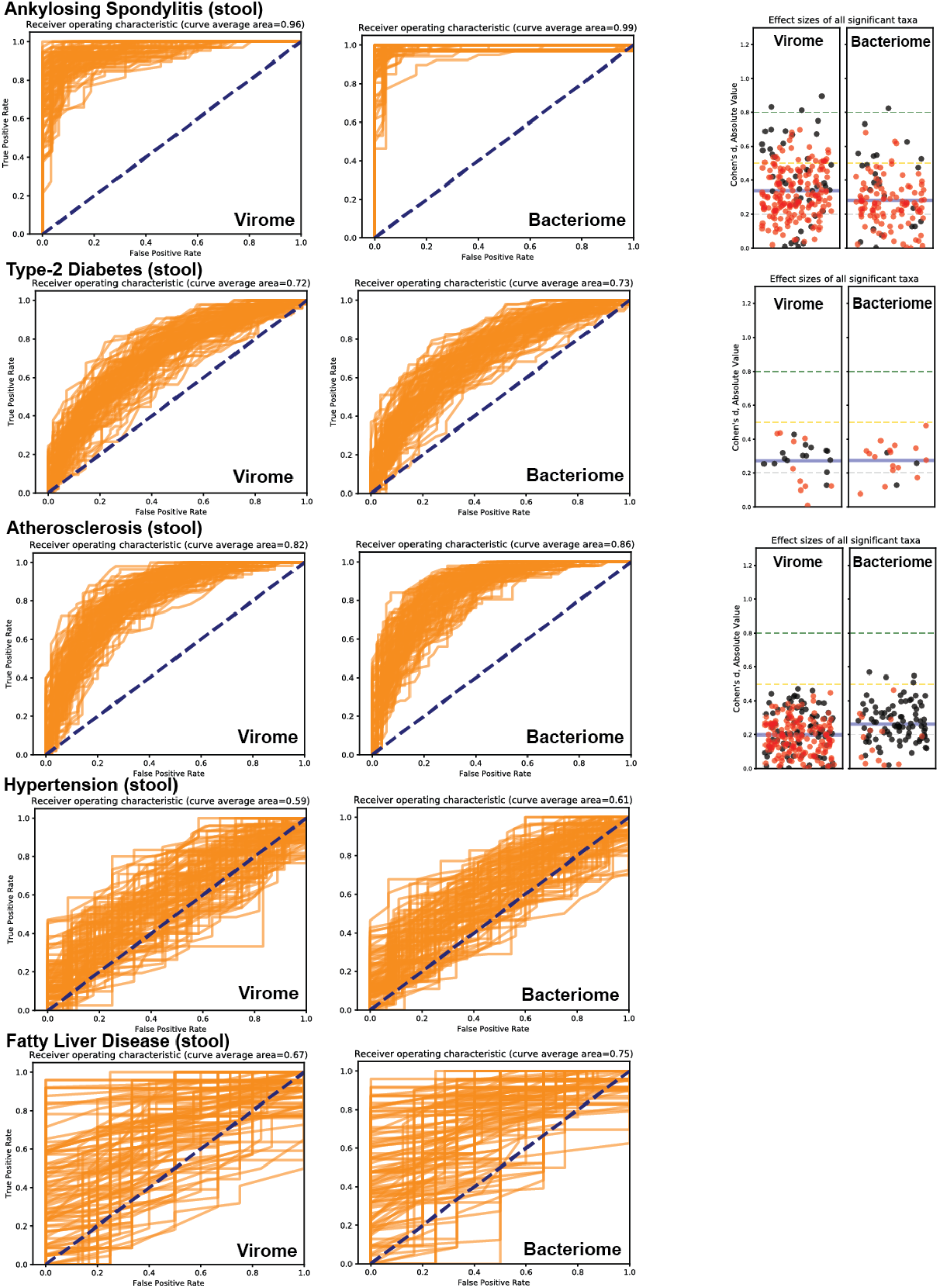
Discriminatory power of viromes and bacteriomes in additional chronic diseases 2. Receiver operating characteristic plots for virome (left panels) and bacteriome (middle panels) data, as described in Figure 6. Summary of effect size data for significant OTUs (right panel), as described in Figure 6.

## Discussion

This study shows that, by leveraging virus-specific hallmark genes, it is possible to mine human metagenomic data at a large scale to create a comprehensive database composed largely of previously unknown virus sequences. This advance, in turn, revealed hidden associations between a variety of chronic disease states and specific virus taxa. It should be stressed that association does not necessarily imply causation, and a variety of associative relationships between viruses and a given disease state are possible. For instance, virus abundance might simply be an epiphenomenon reflecting bacterial host abundance, the human genetics that predispose people to a disease might also provide a more favorable environment for the virus or its bacterial host, the external causes of a disease may create a more favorable environment for the virus, or the virus may contribute to the disease presentation in some way but ultimately does not cause the disease in isolation from other important factors. Verifying the associations we have detected with independent studies of the same diseases in additional populations will be key to understanding the extent to which the findings presented here are generalizable. If the associations are confirmed, it might be possible to experimentally test the causality question by adding or removing phages of interest from gut ecosystems in animal model systems^78^.

A limitation of the case-control studies analyzed here is that they only consisted of a single timepoint for each subject. Virome composition can be noisy, and longitudinal data on individual patients might be more effective for discerning stable viral populations^10^. This problem may have been partly offset by use of large cohort sizes (mostly over 150 total patients). Another limitation is that the analyzed case-control surveys only used DNA WGS methods whereas RNA sequencing of metatranscriptomes might have provided more functional data on expression of specific viral genes, potentially leading to testable hypotheses about possible mechanisms of action. It is also conceivable that correlations for viruses with RNA genomes would be uncovered. Despite these limitations, the current study shows that, using random forest classifiers, the virome may well be more diagnostic than the bacteriome for a variety of chronic diseases. The strong associations of specific virus OTUs in chronic diseases, along with medium to large effect sizes for many OTUs, cries out for more mechanistic investigation of possible causal roles for viruses in chronic human disease.

Even with the relatively inclusive criteria used by Cenote-Taker 2 (discernable amino acid similarity of a viral hallmark gene to a protein the RefSeq virus database), thousands of viruses that live on humans from this dataset could not be taxonomically classified, suggesting that additional families of as-yet-unidentified viruses await formal discovery.

## Methods

### Identification of viral contigs in assemblies

Human Microbiome Project studies and other arbitrarily selected human metagenome studies (Supplemental Table 1) were downloaded from SRA and unique Biosamples were delineated. All runs from a given Biosample were downloaded concurrently, preprocessed with Fastp^79^, and co-assembled with Megahit^80^ using default settings. Subsequent contigs were fed to Cenote-Taker 2, with settings to consider circular or LTR-bearing contigs of at least 1500 nt, ITR-containing contigs of at least 4 kb, and linear contigs of at least 12 kb. These contigs were scanned for genes matching viral hallmark models and terminal repeat-containing contigs with one or more viral hallmark genes were kept, as well as linear contigs with two or more viral hallmark genes. Cenote-Taker 2 hallmark gene database was the September 15th, 2020 version (see GitHub repo: https://github.com/mtisza1/Cenote-Taker2). While Cenote-Taker 2 does take steps to earmark potential plasmids and conjugative transposons, extra precautions were taken by removing ^~^4000 putative viral sequences from the non-redundant database that contained replication-associated but not virion or genome-packaging hallmark genes. For metatranscriptome datasets, all contigs over 1500 nt with RNA virus hallmark genes were kept as putative viral sequences, regardless of end features.

### Clustering similar contigs

Mash^48^ was used to cluster contigs into virus operational taxonomic units (OTUs) due to its ability to handle massive sequence databases, accuracy, and lack of issues arising from genome circularity. All viruses within each higher-level taxon (e.g. *Microviridae*) were used to create Mash sketches (options -k 16 -s 500), and then these sketches were compared to themselves with Mash’s dist function. Within close genomic distances, the value of the Mash distance score is thought to roughly recapitulate average nucleotide divergence. Virus strain-level distinctions are often defined by <5% average nucleotide divergence^51^, so sequence similarity networks were constructed with connections between sequences (nodes) with Mash distance scores ≤0.05 (and p value ≤1e^−10^). MCL clustering^81^ was applied to Mash networks to generate OTU-level clusters. From each cluster of sequences, if circular or ITR-encoding sequences were present, the longest such sequence was used as the representative virus OTU sequence. If only linear sequences were present, the longest linear sequence was used as the representative of the cluster. Singleton contigs (i.e., sequences that were not assigned to any cluster) were also retained for the final database. The same approach was applied for the 99% database (for virus-like particle sequence alignment), but a Mash distance score of ≤0.01 was used.

### Identifying cognate viruses in GenBank and the human Gut Virome Database

Using the NCBI Virus <u>Resource</u>, metadata for all virus genomes listed as complete were downloaded for the following taxa: *Adenoviridae, Anelloviridae, Bromoviridae, Caliciviridae, Circoviridae, Cressdnavircota, Herpesviridae, Luteoviridae, Narnaviridae, Nodaviridae, Papillomaviridae, Polyomaviridae, Tombusviridae, Totiviridae, Tymovirales*, unclassified viruses, unclassified RNA virus, *Virgaviridae*, and all bacteriophage (including prophage). The metadata were sorted so that the longest sequence for each unique species name was selected, and these sequences were subsequently downloaded. Additionally, many GenBank virus genomes simply have a family label followed by the indeterminate abbreviation “sp.” and, as a result, many highly distinct sequences inadvertently share an identical generic label. Therefore, all complete GenBank virus genomes from all non-redundant taxa with an ‘sp.’ designation were downloaded. A Mash sketch was made for the downloaded sequences using options (-k 16 -s 500), and this Mash sketch was compared to the Cenote Human Virome Database Mash sketch (see above). Mash distances of ≤0.05 and p value ≤1e^−10^ were considered to be strict cognate (intraspecies or intrastrain) sequences. Mash distances of ≤0.1 and a p value of ≤1e^−5^ were used for “loose” cognate sequences.

The Gut Virome Database from Gregory et al.^12^ was downloaded from: https://datacommons.cyverse.org/browse/iplant/home/shared/iVirus/Gregory_and_Zablocki_GVD_Jul2020/GVD_Viral_Populations. The same Mash analyses were applied for comparisons with this dataset as with the GenBank database.

### Gene sharing network for unclassified viruses

Vcontact2^52,82^ was run using all RefSeq v88 bacteriophage genomes with recommended settings and all viruses from the Cenote Human Virome Database that were labeled “unclassified” in the taxonomy field. The resulting network was displayed in Cytoscape^83^ and colored manually.

### Virus cores

Using all sequences from Cenote Human Virome Database (i.e. virus OTUs per 95% clustering), virus core coordinates were obtained. The Cenote-Taker 2 hallmark scan outputs for each sequence were used to identify which genes encoded virus hallmarks (i.e. “hallmark genes”). The stop and start coordinates for each hallmark gene were compiled from the Cenote-Taker 2 output amino acid files. The coordinates with the lowest value and highest value were taken, and bioawk was used to trim each fasta nucleotide sequence to start and end with these coordinates, discarding outside sequences.

### Bacteria-encoded CRISPR spacer analysis

CrisprOpenDB (https://github.com/edzuf/CrisprOpenDB, manuscript not currently available for citation) was used (commit 04e4ffcc55d65cf8e13afe55e081b14773a6bb70) to assign phages to hosts based on CRISPR spacer match. Three mismatches were allowed for hits. For hits to bacteria without a currently assigned genus, family-level or order-level taxonomical information was pulled from the output table, when possible.

### Phage-encoded CRISPR spacer analysis

All virus OTU sequences were processed with MinCED (https://github.com/ctSkennerton/minced) to discover CRISPR spacer arrays. As phages can encode CRISPR arrays with spacers as short as 14 nucleotides^84^, MinCED was allowed to detect arrays with spacers of 14 or more nucleotides. The CRISPR-array regions of phage genomes were masked using Bedtools maskfasta^85^, and then all virus OTUs were queried with BLASTN against a database of the CRISPR spacers.

Only hits aligning to the entire length of the spacer and with the following criteria were kept: perfect matches to spacers 16-20 nucleotides, matches to spacers 20-27 nucleotides where (mismatches + gaps) is 1 or 0, matches to spacers ≥ 28 nucleotides where (mismatches + gaps) is 2 or fewer.

### Determining abundance of individual virus OTUs in metagenomes

The final database of “virus core” sequences was processed by RepeatMasker to remove low-complexity regions which recruit reads non-specifically^86^. In the interest measuring known human viruses that may have been missed by Cenote-Taker 2 analysis of human metagenomes, 410 viruses that reportedly infect humans were downloaded using the NCBI Virus Resource, processed by RepeatMasker, and added to the final database for read alignment (full-length sequences were kept for GenBank records). Bowtie2^87^ was used to align reads to the database, and samtools^88^ idxstats was used to calculate read coverage and reads per kilobase per million reads (RPKM) for each contig.

### Comparing OTU abundance and discriminatory ability in case-control studies

For each Bioproject, case vs control samples were determined, if possible, using supplemental table 15 from Nayfach et al^75^, as patients on confounding medications were removed in this table. For other Bioprojects, metadata was taken from SRA^89^ run selector (Supplemental Table 6). For all samples, reads were downloaded from the SRA and trimmed and quality-controlled with Fastp^79^. To quantify abundance of bacterial taxa in each sample, IGGsearch was used with the “--all-species” option.

Wilcoxon rank-sum test was computed with 100 bootstraps using Python, NumPy and SciPy^90^ for each OTU in a given study where at least 10% of the total samples had an RPKM of at least 0.05 (bacterial OTUs with “IGGsearch abundance” of at least 0.005 in at least 10% of the samples were kept). False discovery rate (< 1%) was determined with the Benjamini-Hochberg method using Scipy. Cohen’s d effect size was calculated for each OTU above the significance threshold using DaBest Python package^91^ with 5000 bootstraps.

Random Forest Classifiers from scikit-learn were used^92^. Training/test set sizes were 70%/30%, number of estimators was 100, and a different seed was used for each of the 100 Random Forest Classifiers trained on each dataset.

## Data availability

The fundamental data can be downloaded from: https://zenodo.org/record/4069377. At the time of initial manuscript submission, deposition of each OTU in the CVHD into GenBank is in process and accession numbers will be provided in a future revision.

## Contributions

Michael J. Tisza: Conceptualization, Resources, Data curation, Formal analysis, Validation, Investigation, Visualization, Methodology, Project administration, Writing manuscript Christopher B. Buck: Conceptualization, Resources, Project administration, Editing manuscript

## Acknowledgements

The authors would like to acknowledge Gabriel Starrett and Nathan Fons for their helpful discussions on statistical analyses of the data types used in this paper. This work utilized the computational resources of the NIH HPC Biowulf cluster. (http://hpc.nih.gov).

